# tRNA-derived small RNAs target transposable element transcripts

**DOI:** 10.1101/085043

**Authors:** Sarah Choudury, R. Keith Slotkin, German Martinez

## Abstract

tRNA-derived RNA fragments (tRFs) are 18-26 nucleotide small RNAs that are not random degradation products, but are rather specifically cleaved from mature tRNA transcripts. Abundant in stressed or viral-infected cells, the function and potential targets of tRFs are not known. We identified that in the unstressed wild-type male gamete containing pollen of flowering plants, and analogous reproductive structure in non-flowering plant species, tRFs accumulate to high levels. In the reference plant *Arabidopsis thaliana*, tRFs are cleaved by Dicer-like 1 and incorporated into Argonaute1 (AGO1), akin to a microRNA. We utilized the fact that many plant small RNAs direct cleavage of their target transcripts to demonstrate that where the tRF-AGO1 complex acts to specifically target and cleave endogenous transposable element (TE) mRNAs produced from transcriptionally active TEs. The data presented here demonstrate that tRFs are *bona-fide* regulatory microRNA-like small RNAs involved in the regulation of genome stability through the targeting of TE transcripts.

## INTRODUCTION

The development and refinement of high-throughput small RNA (sRNA) sequencing technologies have allowed the field to progressively uncover layers of non-coding RNA (ncRNA) populations involved in the regulation of diverse processes (1,2). The best-studied sRNAs are microRNAs, which are key molecules governing development and stress responses (3). In addition to microRNAs, some sRNAs derived from ncRNAs such as ribosomal RNAs (rRNAs), small nuclear RNAs (snRNAs), small nucleolar RNAs (snoRNAs) and transfer RNAs (tRNAs) are not random degradation products, but rather have specific processing and functions post-processing (4-6). sRNAs derived from tRNAs of have gained particular attention and interest (7). tRNAs, apart from their primordial role in the central dogma, have been suggested to exert a double regulatory function from being part of a distinct control layer through the production of different sRNAs derived from their transcripts (7). Two different classes of sRNAs derived from tRNAs have been reported, tRNA halves of 30-36 nucleotides (nt) and 18-20 nt tRNA-derived RNA fragments (termed tRFs) (8). tRNA halves inhibit protein translation in a wide range of organisms (9-11),while tRFs have only recently been characterized in depth and their function remains unknown (7). One striking similarity is the accumulation of both tRNA halves and tRFs only in stressed cells (7,12-15).

In mammals, 18-22 nt tRFs may derive from either the 5’, 3’ and tail regions of the mature tRNA transcripts and have been proposed to be both Dicer-dependent (16,17) or -independent (5,6). tRF processing results in the precise generation 18-22 nt sRNAs that are loaded into various Argonaute (AGO) proteins (14,18). These tRF-AGO complexes are biologically active: in human cells they reduce the accumulation of a luciferase reporter construct engineered with a primer binding site site containing a tRF target site (16). Because of their Dicer- and AGO-dependent biogenesis pathway, it has been proposed that tRFs could function akin to a microRNA to inhibit the translation or cleave partially-complementary target sites, however the identification of their endogenous targets has yet to be determined (7).

TE mobilization is detrimental to host fitness as it can cause mutations via insertion, promoter modifications, abnormal genome rearrangements and chromosome breakage (19). In order to restrict their potentially detrimental effects, eukaryotic organisms have evolved multiple mechanisms based on RNA silencing and chromatin modification to control TE expression and maintain TEs in a quiescent silenced state (20). Defeat of host epigenetic silencing and TE transcriptional reactivation occurs upon certain developmental stages, stresses and mutants of various chromatin-modifying proteins (21). One such Arabidopsis mutant is *ddm1*(*DECREASE IN DNA METHYLATION 1*), which lacks the DDM1 *swi/snf* family chromatin remodeler protein (22). TE transcriptional reactivation also takes place naturally in the Arabidopsis pollen vegetative nucleus, which undergoes a process termed developmental relaxation of TE silencing (DRTS) (21) due to the lack of centromeric and heterochromatin condensation in this cell type (23), which may be linked to the lack of DDM1 protein accumulation (24). In both *ddm1* mutants and the pollen grain, transcriptional activation of TEs is results in their degradation into small interfering RNAs as a second-line post-transcriptional defence inhibiting TE activity. RNA is triggered by sRNAs to cleave the TE mRNAs and initiate RNA-dependent RNA Polymerase (RDR) activity to create double-stranded RNA, which is further Diced into abundant secondary siRNAs (24). However, the identification of the primary (non-RDR-dependent) sRNAs that initiate TE RNAi is incomplete.

We utilized the fact that plant sRNAs often cleave their target transcripts to determine the biogenesis pathway and endogenous targets of tRFs. We found that 19 nt tRFs derived from the 5’ region of mature tRNA sequences (tRF-5s) accumulate to high levels specifically in pollen. tRF-5s biogenesis involves the core components of the plant microRNA pathway: Dicer-like 1 (DCL1) and AGO1, and we identify the primary targets of tRFs are TE mRNAs. In addition, we find that the pollen / reproductive tissue-specific accumulation of 19 nt tRF-5s is a characteristic that has been deeply conserved among the plant lineage.

## MATERIAL AND METHODS

### Plant material

Arabidopsis plants of the ecotype Col-0 were grown under standard long day conditions at 22 °C. Inflorescence tissue was used in each experiment unless otherwise noted. Mutant alleles used are described in Supplemental Table 1.

### Transgenic constructs

KRP6-tRF/sRNA target-H2B PCR fragment was cloned as in (25).

### Microscopy Analysis

Pollen grains of T1 plants were mounted in slides containing 50% glycerol and analyzed under a Nikon-A1 fluorescence microscope. Between 4/8 independent transgenic events and 300 and 500 pollen grains per individual transformation event were analysed under the same values of exposure time and fluorescence intensity. GFP intensity in the VN of individual pollen grains was measured through the Nikon-Elements application.

### Small RNA deep sequencing and analysis

Inflorescence sRNAs for library preparation were enriched with the mirVana miRNA Isolation Kit (Life Technologies). The sRNAs for wt Col pollen biological replicate GSM1558871 were not size selected before library production, and thus contain 19nt tRFs. The sRNAs for wt Col pollen biological replicate GSM1558877 and *ago1*pollen GSM1558878 were size selected as in (25). sRNA library production and pre-analysis was done as in (25). Mature tRNA sequences were obtained from the tRNA genomic database (http://gtrnadb.ucsc.edu/). tRNA loci and surrounding regions were extracted from the genome version indicated in the tRNA genomic database. sRNA alignments were performed using PatMaN through the UEA sRNA workbench (26). Predictions of tRF target genes were made using the psRNA-Target (27) using a mRNA/tRF pair scoring cutoff of E ≤ 5. PARE-seq libraries were analyzed with PAREsnip through the UEA sRNA workbench (26). The default high stringency settings were used, removing the t/rRNA filtering step. Protein coding genes and TEs identified through this method were further analyzed by aligning PARE-seq reads to their sequences in order to confirm the predicted target sites. The raw sequencing and genome-matched sRNAs created for this study are available from NCBI GEO repository number GSE63879.

### Total RNA, sRNA northern blot, RT-PCR and 5’RLM RACE-PCR analysis

Total RNA was isolated using TRIzol reagent (Life Technologies). For tRF or microRNA Northern blot detection, 30 or 10 µg of sRNA-enriched RNA were loaded in each lane for inflorescence or pollen Northern blots respectively. Immunoprecipitated sRNA Northern analysis was performed as in (28). sRNA gel electrophoresis, blotting, and cross-linking were performed as described in Pall et al (29). For mature tRNA total RNA and 8% polyacrylamide gels were used. cDNA was generated using an oligo-dT or 3’ region reverse primer and Tetro Reverse Transcriptase (Bioline). Cleavage RACE-PCR was performed as in (28). PCR primers and oligonucleotides used for probe synthesis are listed in Supplemental Table 2.

## RESULTS

### Specific accumulation of 19 nt 5’ derived tRFs in the Arabidopsis pollen grain

In order to characterize ncRNA-derived sRNAs in Arabidopsis, we analyzed the accumulation of ncRNA fragments ranging from 18 to 25 nt in the pollen grain and inflorescence (flower bud)(Figure 1A-B). This analysis revealed that both rRNA and tRNA-derived sRNAs accumulate to high levels in pollen, while, microRNAs decrease in accumulation as previously described (30) (Figure 1A; see Supplemental Figure 1A for analysis of individual bioreplicates). Both sRNAs derived from rRNAs and tRNAs undergo an increase in pollen (rRNA 4.14-fold and tRNA 5.2-fold relative to inflorescence; Figure 1B), although the rRNA-derived sRNAs increase is contributed by all sizes of sRNAs, suggesting potential increased degradation and/or transcription of rRNAs (Figure 1A). In contrast, tRF increase is contributed by a single size class of 19nt sRNAs, suggesting specific processing (Figure 1A and Supplemental Figure 1A). The tRFs size profile ranging from 18 to 25 nt in sRNA libraries from leaf, inflorescence and pollen (from additional 3 bioreps to confirm our initial observation) revealed that 19 nt tRFs specifically accumulate to a higher level in the pollen grain (29 fold higher, Figure 1C, Supplemental Figure 1B). We further validated this result by Northern blot, confirming both the over accumulation of a 19 nt tRF (from the Ala-AGC tRNA) and the lower accumulation of a microRNA (miR161) in pollen (Figure 1D).

**Figure 1.**
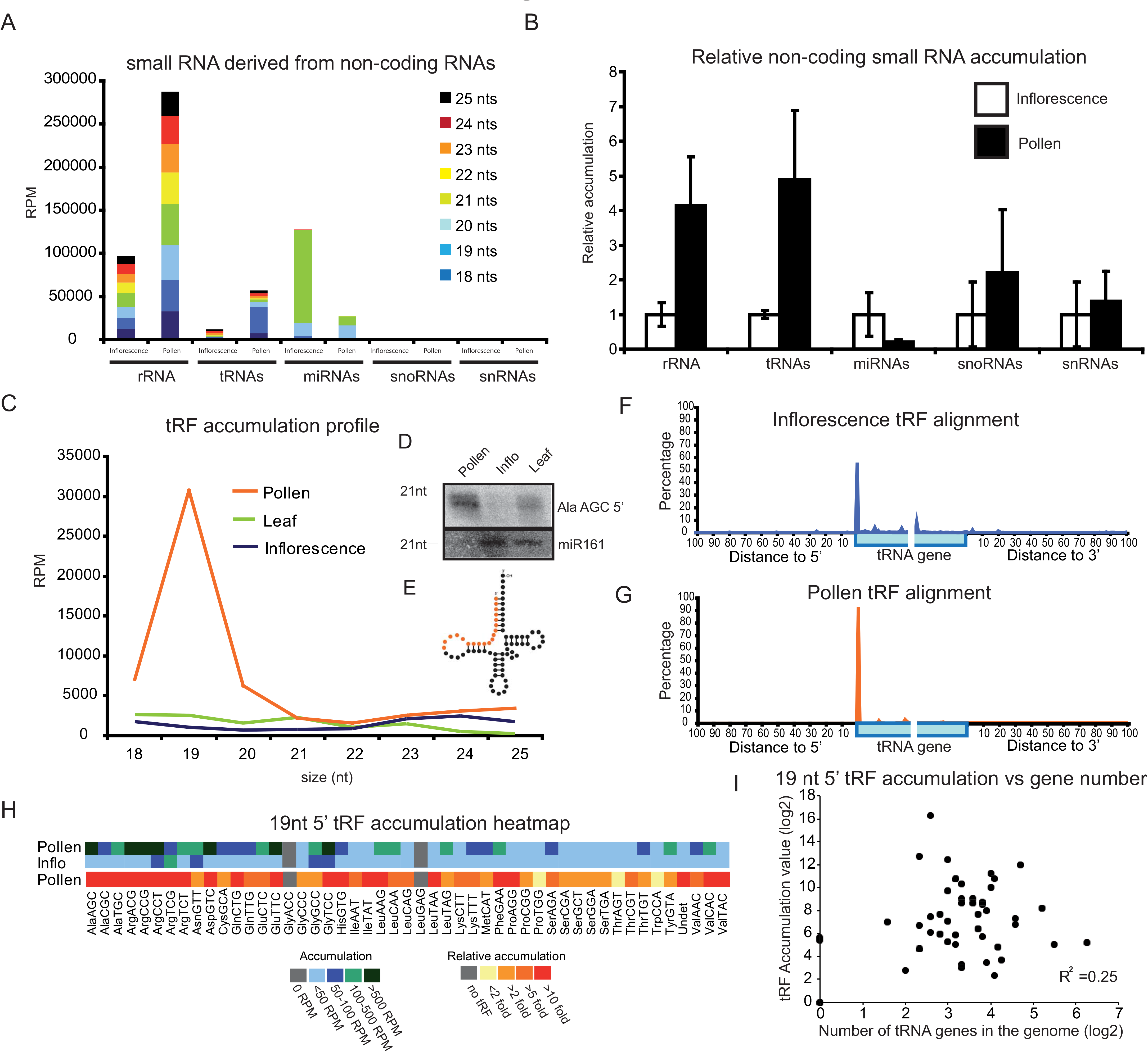
Characterization of tRFs in *Arabidopsis* pollen. **(A)** Analysis of ncRNA-derived sRNAs in inflorescences and pollen. Values are averages of two independent bioreplicates. **(B)** Analysis of the relative accumulation of ncRNA-derived sRNAs in pollen relative to inflorescence (two bioreplicates for each tissue). Error bars represent standard deviation among biological replicates. The inflorescence mean value is normalized to 1.0. **(C)** tRF accumulation profile in different sRNA libraries derived from leaf (3 bioreplicates), inflorescence (2 bioreplicates) and pollen (5 bioreplicates). **(D)** Northern blot detection of 18-19 nt AlaAGC 5^’^-derived tRF and miR161 in pollen, inflorescence and leaf. **(E)** Cartoon representation of tRNA secondary structure with nucleotides that give rise to tRF-5s coloured in orange. **(F-G)** Distribution of 5^’^ end nucleotide of 19 nt sRNAs over all tRNAs and a 100 nt window both 5^’^ and 3^’^ of the tRNA gene in inflorescence (F) and pollen (G). **(H)** Total RPM and fold change heatmaps of 19 nt 5^’^-derived tRF accumulation in pollen relative to inflorescence. **(I)** Scatterplot of the log2 normalized tRF accumulation level in the pollen grain for each individual tRNA gene against the log2 number of tRNA genes present in the Arabidopsis genome.

tRFs can derive from both the 5^’^ and the 3^’^ ends of the mature tRNAs (7). In order to determine the processing features of tRF over-accumulation in pollen, we plotted 19 nt tRFs along tRNA genes and 100 nt on either side of each mature tRNA coding region. We found that tRFs accumulate from the 5^’^ end of the mature tRNA (Figure 1D-G). In inflorescence tissue, where few tRFs accumulate (Figure 1C), there is a preference for 5^’^ processing of the mature tRNA transcript into tRFs (49%, Figure 1F), while in pollen this 5^’^ processing preference is increased (88%, Figure 1G) indicating that in pollen 5^’^-derived tRF biogenesis is more specific and/or efficient. We further checked if tRF specific processing from the 5^’^ region is masked by the use of tRNA genomic sequences lacking the CCA sequence that is added to the 3^’^ end of tRNAs post-transcriptionally (31), and this analysis revealed that tRFs are only processed from the 5^’^ end of mature tRNA sequences (termed tRF-5s, Supplemental Figure 1C-D). From the 49 tRNAs grouped by codon type in the Arabidopsis genome, tRFs accumulate for 47 (95.92%, Figure 1H). Intron presence (SerGCT, MetCAT and TyrGTA), or the number of genes coding for each specific tRNA codon type in the genome, did not influence tRF-5s accumulation (Figure 1I). Together, our data suggests that tRF formation is a general phenomena for all tRNAs in Arabidopsis pollen.

To determine if tRF formation in pollen is found in other plants known to accumulate tRFs in somatic tissues (32), we investigated published data in rice and maize. We found that the specific 5^’^ processing of 19 nt tRF-5s in pollen was conserved in rice and maize (Figures 2A-B, 2D-E, 2G-H and Supplemental Figure 2). In order to analyse if the phenomena was a characteristic potentially conserved also in non-flowering land plant precursors, we analysed the tRF distribution in the reproductive tissue of the bryophyte *Physcomitrella patens*. The Physcomitrella life cycle comprises an alteration of two generations. First, a haploid gametophyte (a filamentous structure termed protonema) derived from spores that produce plant-like structures from their buds (termed gametophore). These gametophores give rise to the gametes which after fusion will produce a diploid sporophyte (33). The analysis of the sRNA libraries available from gametophore-sporophyte compared to protonemata and protonemata-young gametophore revealed that non-vascular plant precursors accumulate 5^’^-derived 19 nt tRFs from their reproductive structures (Figures 2C, 2F and 2G-H). Together, these results reveal that in plants, tRFs undergo specific processing from the 5^’^ region of the mature tRNA transcript (tRF-5s) in pollen or analogous male gametophyte structure.

**Figure 2.**
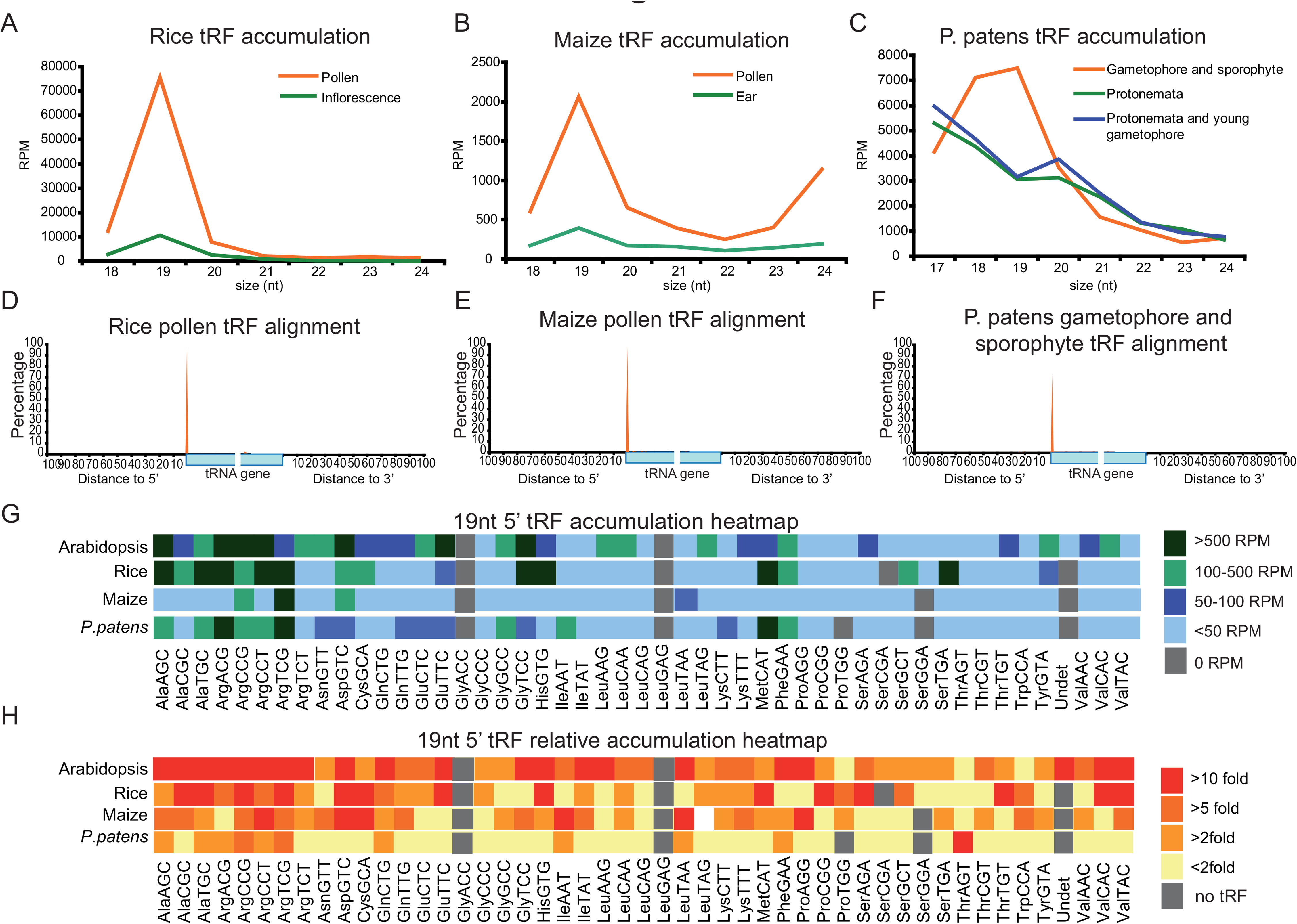
Accumulation of 19 nt 5^’^-derived tRFs is conserved in land plants. **(A-C)** tRF accumulation profile in pollen and leaf from rice (A) and maize (B), and from gametophore-sporophyte, protonemata and young gametophore-protonometa from *Physcomitrella patens* (C). **(D-F)** Distribution of 19 nt tRFs along tRNA loci in pollen from rice (D) and maize (E), or gametophore-sporophyte from *Physcomitrella patens* (F). **(G-H)** Total RPM (G) and fold-change (relative accumulation of 19 nt tRF-5s in reproductive compare to somatic structures) heatmaps (H) of 5^’^-derived 19 nt tRFs of Arabidopsis, rice and maize pollen and *Physcomitrella patens* gametophore-sporophyte.

### *ddm1* mutants display tRF accumulation

The loss of heterochromatin and activation of TEs in wild-type pollen is mimicked in *ddm1* mutant plants (24). This led us to explore whether *ddm1* mutants have increased tRF accumulation in the somatic plant body. We found 19nt tRFs also accumulate to higher levels in inflorescence tissue of *ddm1* mutants (4.04 mean fold increase, Figure 3A), although this reactivation is lower and more variable between bioreplicates (Supplemental Figure 3A). This accumulation is not related with earlier development of the anthers in *ddm1* mutants, as 19 nt tRFs also over-accumulate in *ddm1* seedlings (Supplemental Figure 3B). The 5^’^ preferential processing of tRFs in pollen was also maintained in *ddm1* inflorescences (Figure 3B and Supplemental Figure 3C). We further confirmed by Northern blot the higher level of accumulation for 19 nt tRF-5s, indicating the validity of our sRNA high-throughput sequencing analysis (Figure 3C). Although the distribution of 19 nt tRF over-accumulation in *ddm1* was similar to the one experienced in pollen, fewer tRNAs increase tRF accumulation in *ddm1* compared to wt Col inflorescences (Figure 3D).

**Figure 3.**
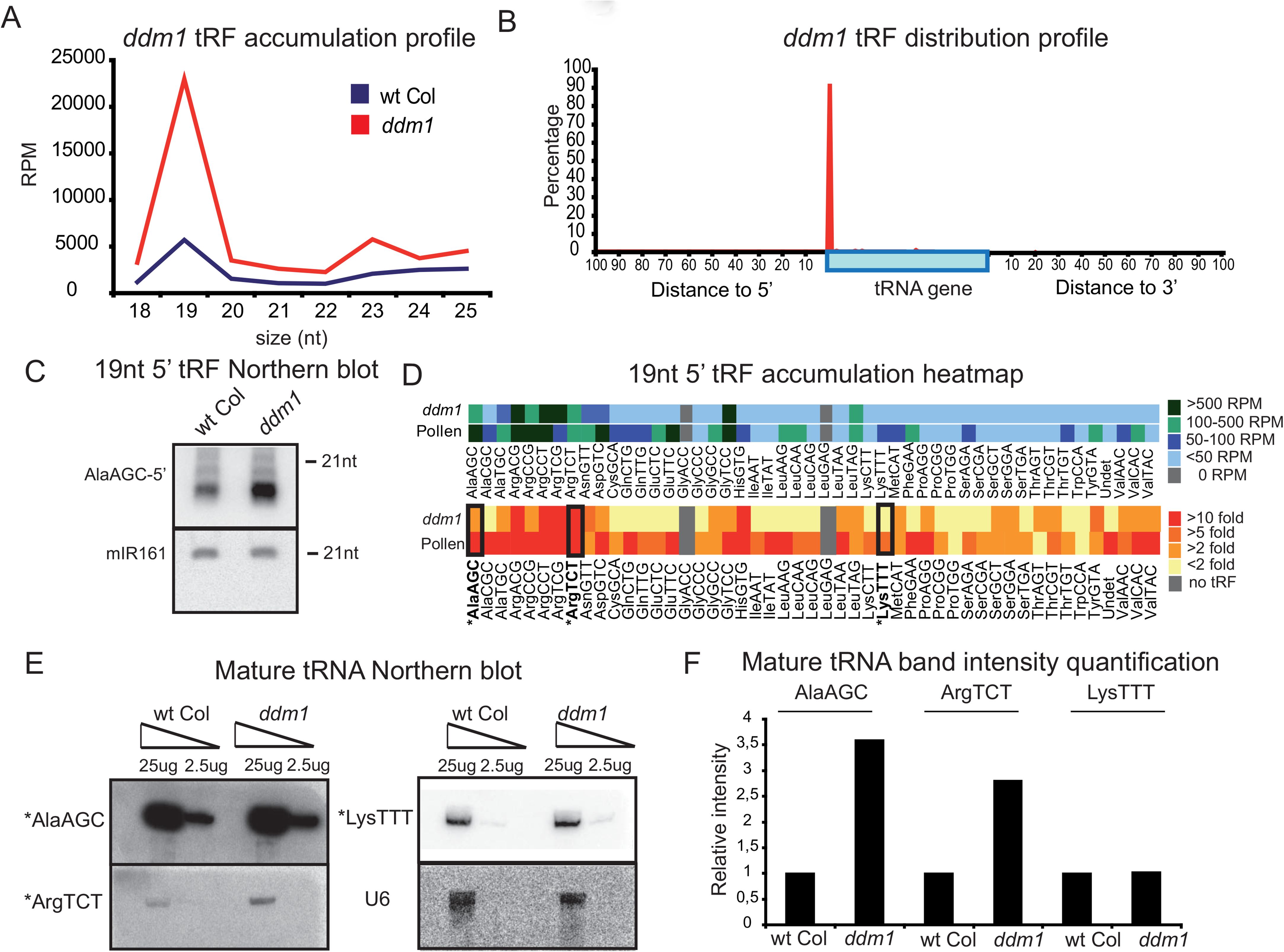
DDM1 influences the accumulation of tRFs. **(A)** tRF accumulation profile in *ddm1* compared to wt Col inflorescence tissue. The average value of 3 bioreplicates is shown. **(B)** Distribution of 5^’^ end nucleotide of 19 nt tRFs on tRNA loci in *ddm1* inflorescence. **(C)** Northern blot detection of AlaAGC 19 nt tRF-5 in wt Col and *ddm1* backgrounds. miR161 was used as loading control.**(D)** Total RPM and fold change heatmaps of 19 nt 5^’^-derived tRF accumulation in *ddm1* inflorescences and wt Col pollen both relative to wt Col inflorescence. tRNAs used for normalization in panel E are highlighted and marked in bold. **(E)** Northern blot detection of selected mature tRNAs (highlighted in panel D) in total RNA from wt Col and *ddm1*. U6 RNA was used as a RNA loading control.**(F)** Band intensity quantification in *ddm1* relative to wt Col normalized to the U6 intensity for the Northern blots represented in E.

Interestingly, we found that increased levels of the corresponding mature tRNA could explain some tRF accumulation in *ddm1*, while for other tRNAs this correlation did not exist (Figure 3E-F). This suggests that some mature tRNA accumulation levels are under the control of DDM1. Interestingly other mutants with decreased DNA methylation (*met1* and the triple mutant *drm1/drm2/cmt3*,termed *ddc*) also have slightly higher accumulation levels of tRFs (Supplemental Figure 3D). Therefore, both mature tRNA levels and tRF accumulation are influenced by epigenetic regulation, and the known reprogramming of DNA methylation in the pollen grain may contribute to tRF accumulation in this tissue.

### tRFs are processed by the microRNA pathway

Several studies have shown that human tRF biogenesis depends on the miRNA pathway (17,34), although other authors point that DICER could be dispensable for tRF biogenesis (6,32). We find that when tRF-5s accumulate to high levels (in *ddm1* and wt pollen) they display similar processing precision values compared to microRNAs, suggesting that tRF-5s were cleaved with a high precision (Figure 4A). The observed increased precise processing of 19 nt tRF-5s in *ddm1* and pollen led us to examine tRF biogenesis in those tissues. We examined Dicer-like1 (DCL1) due to its known role in microRNA biogenesis (35). We observe reduced tRF accumulation in a *ddm1*/*dcl1*double mutant (Figure 4B and Supplemental Figure 4A), demonstrating that the biogenesis of tRFs is similar to microRNAs (36). We moreover confirmed the accumulation of tRFs by Northern blot in pollen grains from *dcl1* and *ago1* single mutants (Figure 4C). In a similar manner to miRNAs, a mutation in AGO1 does not impairs tRF-5s accumulation (36), suggesting that AGO1 acts downstream of DCL1 and a potential redundancy with other AGO proteins.

**Figure 4.**
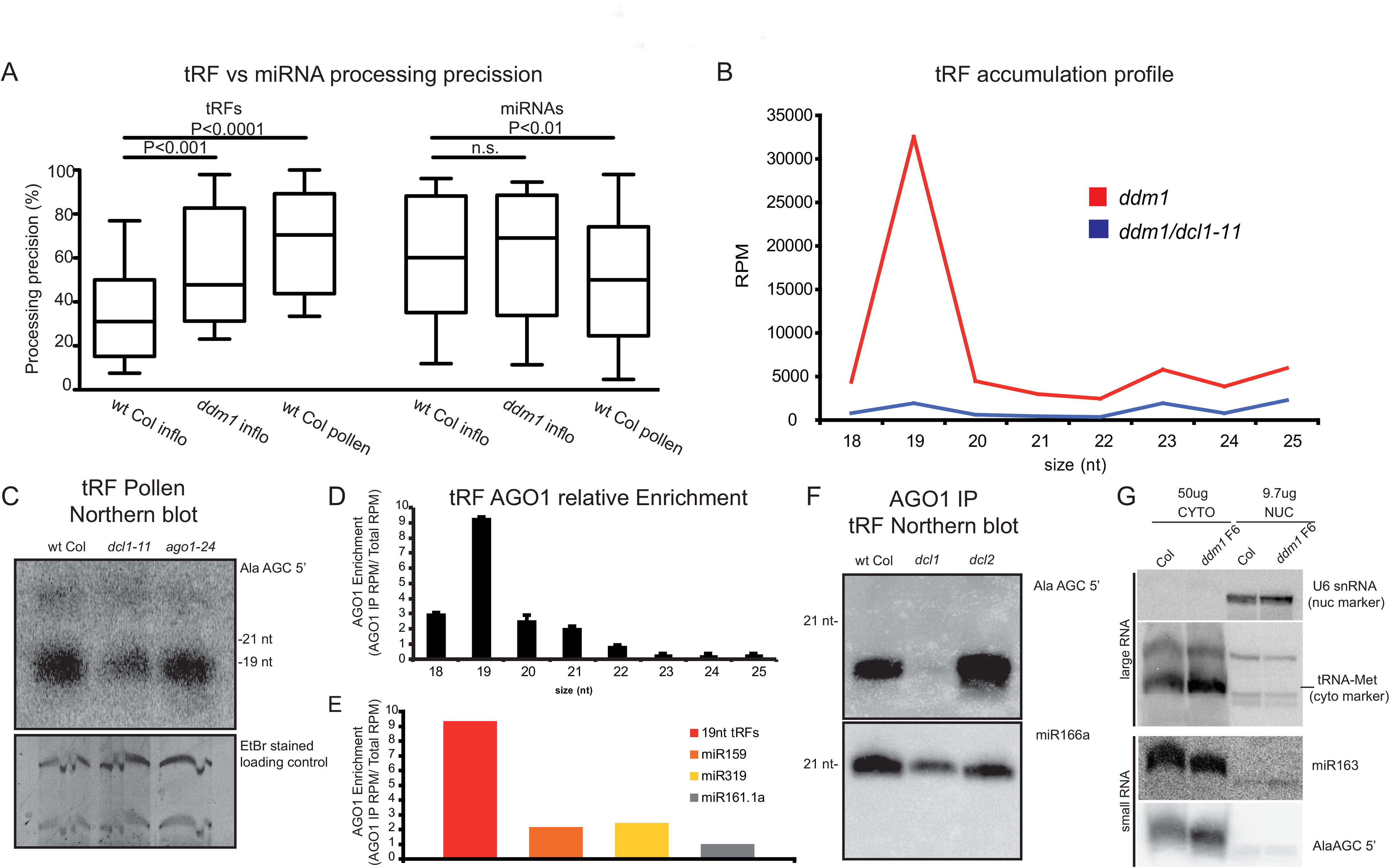
5^’^-derived tRFs have a microRNA-like biogenesis pathway. **(A)** Processing precision of tRFs and microRNAs in wt Col inflorescence, *ddm1* inflorescence and wt Col pollen sRNA libraries. Bars between samples indicate the P value as determined through Student t-tests (two tailed, 95% of confidence interval). n.s. = non-significant difference. **(B)** tRF accumulation profile in *ddm1* and *ddm1/dcl1* **(C)** Northern blot detection of 19 nt AlaAGC tRF-5 in wt, *dcl1* and *ago1* pollen grains. **(D)** tRF enrichment profile in wt AGO1 immunoprecipitated sRNAs relative to accumulation in inflorescences. **(E)** Fold enrichment of tRFs and selected AGO1-loaded miRNAs or non-AGO1 loaded (miR161.1a) **(F)** Northern blot detection of AlaAGC 5^’^ 19 nt tRF in AGO1 immunoprecipitated sRNAs from wt Col, *dcl1* and *dcl2*. **(G)** Northern blot detection of 19 nt AlaAGC tRF-5 in RNA extracted from cytoplasmic and nuclear fractions. miR163 and the mature form of tRNA Met were used as cytoplasmic RNA controls and the snRNA U6 was used as a control of nuclear RNA.

Previous bioinformatic analysis of public sRNA libraries reported that 19 nt tRFs are loaded preferentially into AGO1 in wt Col-0 Arabidopsis (32,37). We next explored whether tRFs are incorporated into AGO1 by analyzing sRNA libraries constructed from AGO1 immunoprecipitates derived from wt Col and *ddm1* inflorescences (Figure 4D and Supplemental Figures 4B-C). In our AGO1-immunoprecipitated sRNA libraries, 19 nt tRFs are enriched 9-fold relative to wt inflorescences (Figure 4D), higher than some miRNAs (Figure 4E). In fact, tRFs represented a substantial portion of AGO1-immunoprecipitated sRNAs: 11.24% (wt) and a 17.14% (*ddm1* background) of the ncRNA sRNAs (Supplemental Figure 4C). In addition, we verified the presence of tRFs in AGO1 through Northern blot detection of AGO1-immunoprecipitated sRNAs (Figure 4F). The lack of accumulation of this tRF in AGO1-immunoprecipitated sRNAs from *dcl1* mutant plants indicates the specific AGO1 loading of DCL1 processed tRFs (Figure 4F).

Next, we analyzed the cellular localization of tRFs by Northern blot in RNA extracted separately from nuclear and cytoplasmic fractions. 19 nt tRF-5s have a cytoplasmic accumulation pattern (Figure 4G), similar to that of AGO1-loaded microRNAs. In addition, we further confirmed the lack of polyadenylated tRNA transcripts, which could potentially feed into the miRNA biogenesis pathway, through oligo dT-primed RT-PCR in *ddm1,ddm1/dcl1* and *ddm1/ago1* backgrounds (Supplemental Figure 4D). This data demonstrates that the precursor molecules to tRF-5s are not Pol II-derived transcripts, but rather are mature tRNAs processed by the miRNA pathway (Supplemenatal Figure 4D).

### tRFs target transposable elements

The loading of tRF-5s into AGO1 led us to question if these sRNAs are active components of the RNA silencing network. First, we performed a bioinformatic prediction of potential tRF target-mRNAs, which showed that the majority of these were protein coding genes (78.82%) and TEs (190, 17.72%) (Figure 5A). In order to check if any of these predicted targets were biologically real, we analyzed the correlation between the predicted 19 nt tRF-5s and mRNA cleavage data from publically available PARE libraries from wt Col inflorescences in a 200 nt window (Supplemental table 3). Plant microRNAs often cleave their target transcripts between the 10/11 nt of the guide microRNA, providing a mRNA cleavage footprint detectable by PARE-sequencing (38). While no obvious preferential accumulation of PARE reads was detected at the predicted target sites for protein coding genes (Figure 5B, 0.98% of PARE reads at the predicted tRF target site), when the same analysis was carried out for TE sequences, an 8.95-fold increase of TE PARE reads was detected at the tRF target sites (Figure 5C). Interestingly, in *ddm1* mutants where both TE mRNAs and tRFs accumulate to higher levels, targeting of TEs by tRFs was amplified 22.9-fold (10 nt surrounding the predicted target site, Figure 5D). This increased targeting in *ddm1* mutants is exclusive to TEs, as protein-coding gene targeting remains low in *ddm1* mutants (Supplemental Figure 5).

**Figure 5.**
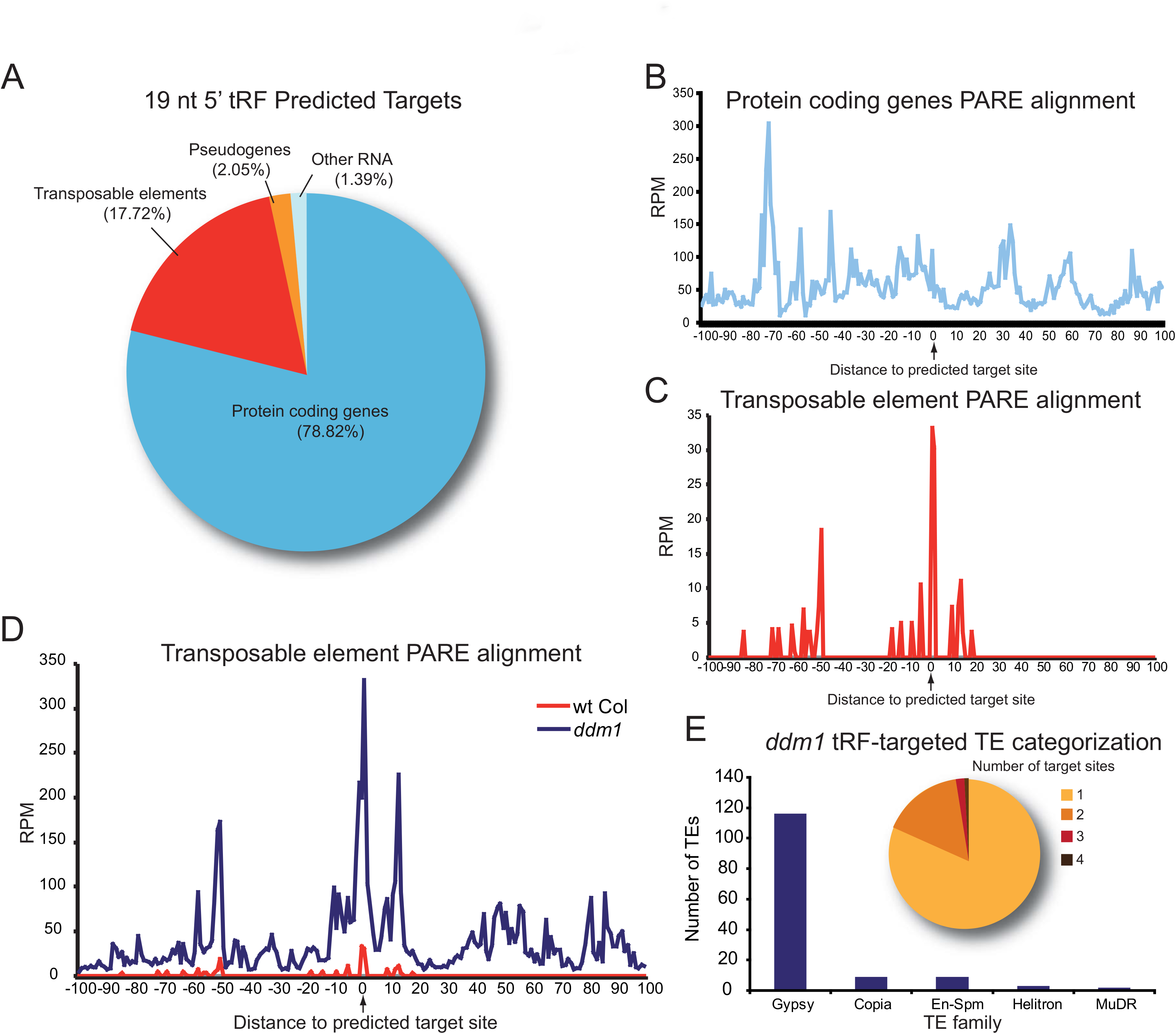
tRFs target transposable elements. **(A)** Categorization of predicted tRF mRNA-targets. **(B-C)** wt Col inflorescence PARE reads alignment along a 100 nt window 5^’^ and 3^’^ to the predicted target sites for protein coding genes (B) or TEs (C). **(D)** PARE reads alignment along a 100 nt window 5^’^ and 3^’^ to the predicted target sites for TEs in wt and *ddm1*. **(E)** *ddm1* tRF-targeted TEs super-family categorization (histogram) and number of tRF target sites for tRF-targeted TEs in *ddm1* (pie chart).

A majority of tRFs in the *ddm1* background targeted TEs belonging to the *Gypsy* family (83.45%, Figure 5E) and had a single tRF target site (81.74%) (Figure 5E). Therefore, our data suggests that when TEs are transcriptionally activated and produce mRNAs, tRFs are able to direct the cleavage of these TE mRNAs.

### Validation of tRF-mediated cleavage of TE mRNAs

To confirm the cleavage ability of tRFs, we selected a 19 nt tRF-5s derived from MetCAT, which accumulates to intermediate levels (Figure 1H) in wt Col-0 and was predicted to target a member of the *Gypsy* TE family, *Athila6A*. We analyzed the specific *Athila6A* cleavage through 5^’^ RLM RACE PCR and detected cleavage only in the *ddm1* background with a high specificity for the predicted tRF cleavage site (9 out of 10 clones sequenced, Figure 6A). Remarkably, this target site is different from the primer binding site for tRNAs that is present in retrotransposons as it is located in the 3^’^ region of the *Athila* element (which is moreover targeted by the 3^’^ region of the mature tRNA). In accordance with tRF-5s biogenesis, this cleavage was dependent on DCL1 and AGO1 (Figure 6B). We next generated STTM lines (39) in the the *ddm1* background (therefore with transcriptionally active TEs) in order to specifically sequester and inhibit the action of the MetCAT 19 nt tRF-5s using as control another transgenic line sequestering a 19 nt tRF-5 derived from AlaAGC. Analysis of the 5^’^RACE product (Figure 6C) and *Athila6A* transcript accumulation level (Figure 6D) indicated that the targeting of the TE by the tRF was abolished in those transgenic lines sequestering 19 nt MetCAT tRF-5.

**Figure 6.**
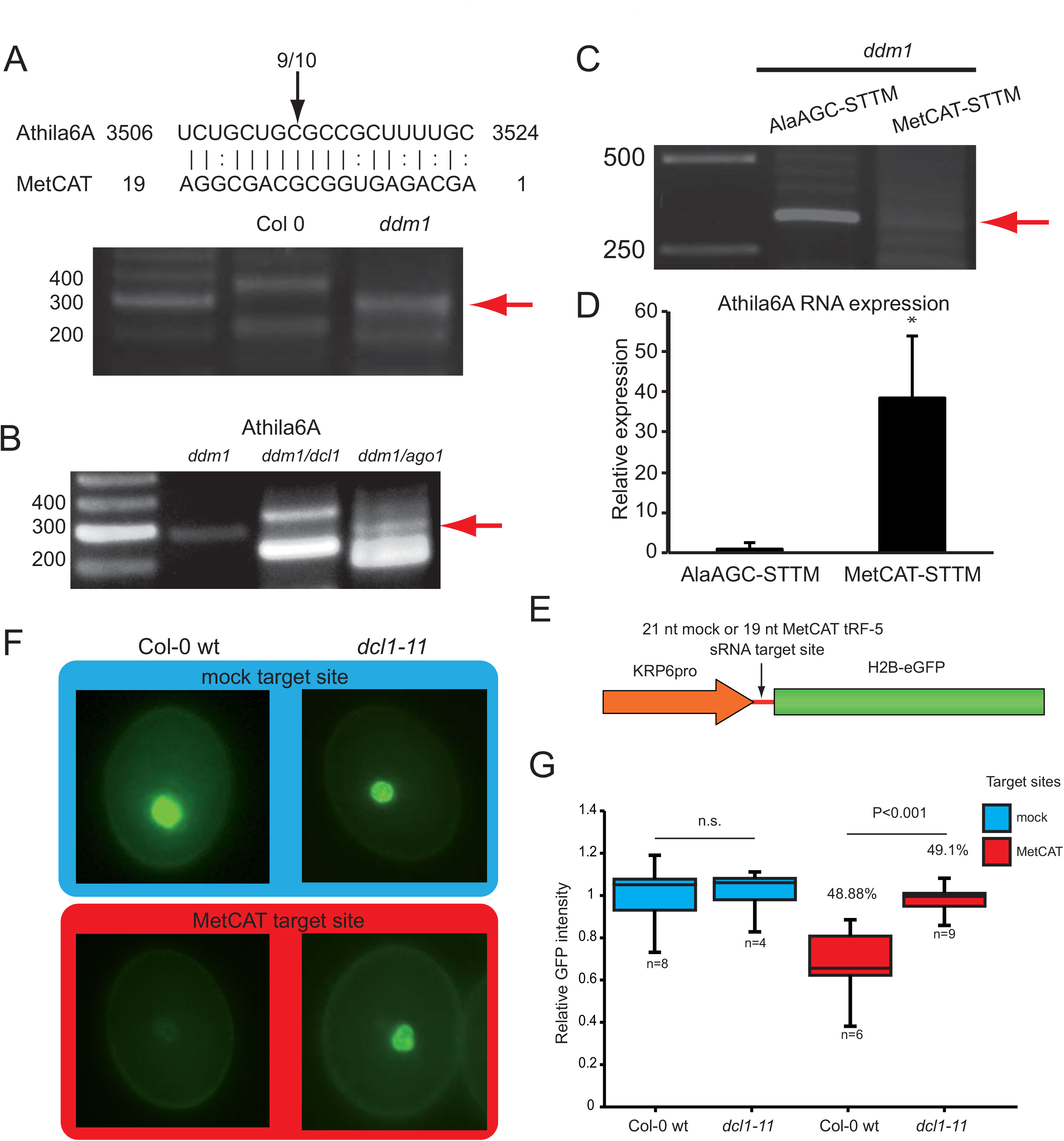
tRF targeting validation. **(A)** tRF alignment with *Athila6A*. Lines indicate prefect complementarity, and dotted interactions specify imperfect non-canonical base pairing sites. Cleavage site and frequency of observed/number of 5^’^RLM RACE PCR product sequences cloned are indicated over the black arrow. The red arrow indicates the cloned band of the expected size that was sequenced. **(B)** 5^’^ RLM RACE PCR cleavage site from *Athila6A* in the *ddm1,ddm1/dcl1* and *ddm1/ago1* background. **(C)** 5^’^ RLM RACE PCR cleavage site from *Athila6A* in SSTM transgenic lines for the AlaAGC and MetCAT 19 nts tRF-5s. 19 nt Ala AGC is not predicted to target *Athila6A.*(**D**) qRT-PCR of *Athila6a* in the same lines as in E with primers spanning the cleavage site. **(E)** Cartoon representation of the transgene constructs used. **(F)** Representative pictures of GFP intensity in pollen grains from wt Col or *dcl1* plants carrying the transgenes from panel D. **(G)** Box plot representation of relative GFP intensity values in wt Col or *dcl1* pollen grains from transgenic lines expressing a construct with a ^“^mock^”^ or MetCAT target site. Values over the boxes represent the percentage of pollen grains with detectable GFP (equivalent to segregation) in the T1. Bars between samples indicate the P value as determined by Student t-test (two tailed, 95% of confidence interval).

We next determined if tRF-5s are functional in the pollen grain, where they accumulate to high levels (Figure 1C). We created a pollen GFP reporter line with a 5^’^UTR containing a 19 nt stretch complementary to the MetCAT tRF-5 or a 21nt ^‘^mock^’^ site not targeted by any endogenous siRNA (Figure 6E), and obtained transgenic lines in both wt and *dcl1* backgrounds. Analysis of GFP fluorescence indicated that the addition of the 19 nt MetCAT tRF-5 complementary sequence resulted in a statistically significant reduction in pollen fluorescence compared to the control target sequence, while when transformed into the *dcl1* mutant background, this difference was abolished (Figure 6E-F). Together, our data confirms the targeting of TE transcripts by tRNA-derived tRFs.

## DISCUSSION

In just the last 7 years new regulatory functions have been discovered for tRNAs, highlighting our lack of complete knowledge of regulatory RNAs and new surprising roles for one of the most well-studied RNA types (40). Here, we have demonstrated that tRNAs are specifically cleaved in plants into 19nt tRF-5s and that these fragments have miRNA-like biogenesis, localization and function (see model in Supplemental Figure 6). Although tRF association with RNA silencing has been demonstrated (6), here we present evidence of at least one additional biological roles. We^’^ve demonstrated that tRFs are part of a genomic protection mechanism through the targeting and cleavage of TE mRNAs, although how this system evolved remains unknown.

Interestingly, several recent papers have linked TEs and tRNAs. Mutations in a subunit of RNaseP (involved in the processing of the 5^’^trailer of tRNA precursors) in Drosophila leads to sterility due to activation of DNA damage checkpoint proteins and TE reactivation coupled with piRNA source collapse (41). Moreover, two miRNAs (miR1511 and miR845, known to target LTR retrotransposons) (42) have precursors that have an evolutionary origin in the tRNA MetCAT (43). Interestingly, here we found that the tRNA MetCAT by itself produces a 19 nt tRF-5 that directly targets the LTR retrotransposon *Athila6A*. tRF production and TE activity seem to have parallel regulation, as both tRFs and TEs are known be expressed and accumulate during stress (12,13,44). In addition, both tRFs and TEs mRNAs accumulate in situations with de-regulated heterochromatin such as *ddm1* mutants and in wt pollen. TE expression is a common phenomenon in wt pollen across flowering plant species (24,45), and we find that tRF accumulation parallels this accumulation in male gametophytes. Interestingly, tRF and tRNA accumulation may be linked to coordinate translation rates. This occurs in human cells (46) and pollen grain germination is dependent on the delayed translation of transcripts accumulated at maturation (47). An access of mature tRNAs awaiting translation to restart could lead to funnelling of tRNAs in to the miRNA pathway (48). Together, our data suggests a conserved natural tRF-5 biogenesis in the male gametophyte of plants and a potential role in gamete protection against intrinsic gametophyte-associated TE reactivation.

## ACKNOWLEDGEMENT

The authors thank Jay Hollick, Jane Jackman and Andrea McCue for helpful comments, and Andrea McCue & Chris DeFraia for their data contributions.

